# A quantitative interaction between signal detection in attention and reward/aversion behavior

**DOI:** 10.1101/032912

**Authors:** V Viswanathan, BW Kim, JP Sheppard, H Ying, K Raman, MJ Lee, S Lee, F Mulhern, M Block, B Calder, D Mortensen, AJ Blood, HC Breiter, Phenotype Genotype Project in Addiction and Mood Disorders

## Abstract

This study examines how processes such as reward/aversion and attention, which are often studied as independent processes, in fact interact at a systems level. We operationalize attention with a continuous performance task and variables from signal detection theory, and reward/aversion with a keypress task using variables from relative preference theory. We find that while the relationship between reward/aversion and attention is functionally invariant, a power law formulation akin to the Cobb-Douglas production function in economics provides the best model fit and theoretical explanation for the interaction. These results indicate that a decreasing signal-to-noise with signal detection results in higher loss aversion. Furthermore, the estimated exponents for the multiplicative power law suggest capacity constraints to processing for attention and reward/aversion. These results demonstrate a systemic interaction of attention and reward/aversion across subjects, with a quantitative schema raising the hypothesis that mechanistic inference may be possible at the level of behavior alone.

## Introduction

Reward/aversion processing and attention have been independently studied for more than 60 years, with quantitative descriptions of matching in reinforcement theory [1,2] and signal detection theory (SDT; [3–5]) providing early examples of mathematical specifications for these behavioral processes. Since these respective frameworks for reward/aversion and attention were developed, they have been extended by development of hedonic deficit theory [6,7], prospect theory [8,9], and relative preference theory [10,11] for reward/aversion, and digital communication incorporating demodulators for attention [12]. Across these reward/aversion frameworks, a basic calibration is described between (a) the intensity and valence of an individual’s emotion and/or wanting, and (b1) an environmental contingency for a goal-object (e.g., the probability of receiving a particular concentration of glucose in a water droplet; [1,2]), (b2) a hedonic deficit around such a goal-object [6,7], (b3) market or group evaluations of a goal-object [8,9], or (b4) the pattern of experience around such goal-objects [10,11]. Of these four frameworks for calibrating value, only one, namely matching, has been related to signal detection during attention [13], and this relationship was (i) purely theoretical and (ii) not validated by neuroscience data or human behavioral data.

Modern overview models of attention include components such as “salience filtering”, “sensitivity control”, and “competitive selection” [14], which relate to aspects of reward/aversion [1,2,6,7,9,11]. Neuroscience data suggests that similar brain regions and neurotransmitters such as dopamine are involved with both reward/aversion and attention [15]. Psychopathology studies report alterations in attention with presumed disorders of reward such as major depression or addiction, whereas addictive substances such as amphetamine can be used to treat attention deficit disorder [16,17,18]. Such results have led some to note that alteration in either reward or attention variables might be used to explain effects attributed to the other [19].

Given this background, we explored if there was a mathematical relationship between reward/aversion and attention variables. We assessed this question in individuals who completed unrelated reward/aversion and attention tasks. We used two independent tasks so that the existence of a mathematical structure could not be predicted from the use of one task, or use of the same experimental stimuli across both tasks. For this proof of existence study, we focused on mathematical formulations of reward/aversion and attention with strong face validity, namely, relative preference theory (RPT) and the original SDT [3,4]. The results of this work have been presented in several talks/meetings previously [e.g., 20,21,22], as the manuscript describing these results has gone through many rounds of submission since early 2013.

RPT models approach and avoidance behavior within an intrinsic motivation-like framework in which no external rewards are provided [23,24], yet participants can produce variable amounts of work [25,26] to modulate the time of stimulus viewing. RPT encodes fundamental features of the other reward theories [11] and is the only formulation of reward/aversion using an information theory variable [27]. To quantify RPT, we used a validated keypress task (e.g., [11,28–35]) with a beauty stimulus set [28] to produce variables K and H that encode, respectively, mean keypress number and Shannon entropy (i.e., information; [27]; Figure 1a). These variables produce a valuation graph, which has been interpreted to relate “wanting” of stimuli [28] to the uncertainty associated with making a choice [11], and closely resembles the value function in prospect theory [8]. Keypress measures of value can be connected to neural systems [28–34]. It should be noted that pattern variables such as the Shannon entropy have been shown to be important metrics for quantifying neural processing [36–38], and define the “information” that is processed in cognitive neuroscience [11,39].

**Figure 1:**
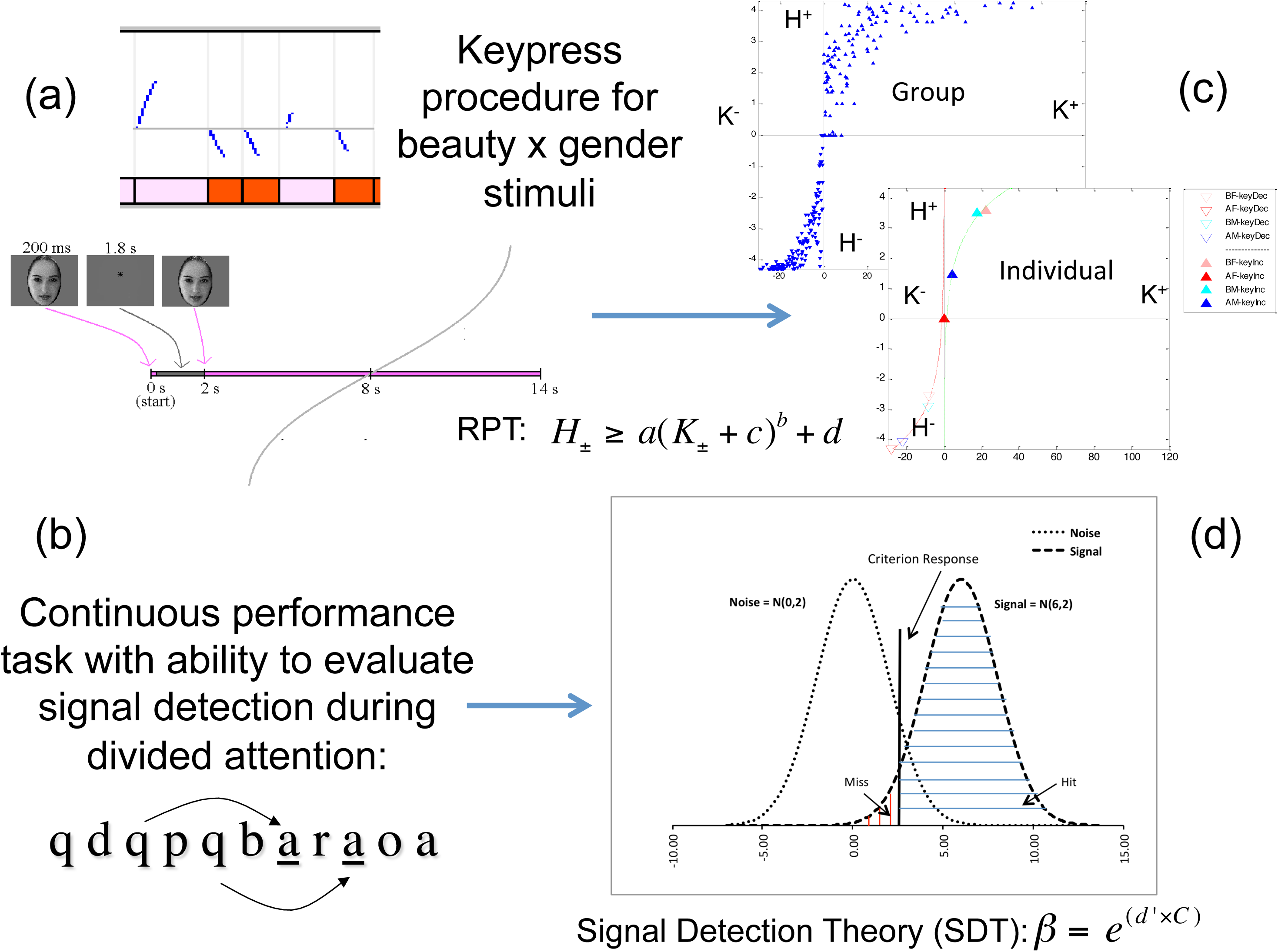
Experimental paradigms. (a) A schema for the keypress paradigm shows, at top, raster-plots of keypress effects on viewing time as blue curves going up or down from a default viewing time of 6 seconds. Pink and red blocks represent presentation of beautiful and average female faces, respectively. Below the raster-plots, timing of face presentation is schematized (see Methods). (b) Results of keypressing produced a boundary envelope for group data and curve fits for individual subjects, for the mean keypress variable K and Shannon entropy H [i.e., information]. (c) A visual continuous performance task quantified signal detection parameters during divided attention. One letter was shown per second in the center of the visual field. Subjects responded when a target letter (“a”) appeared exactly four letters after the cue (“q”). Cue-target pairs could be interleaved, necessitating divided attention. (d) Signal detection analysis allowed quantitation of a criterion response and relative signal-noise distributions for β and d’ variables.

For attention function that could be quantified by SDT we used a continuous performance task (CPT) given it is one of the most widely used neuropsychological measures of attention [40–42]. Subjects participating in a CPT view a continuous presentation of changing stimuli, such as a sequential presentation of letters, and respond to designated targets. Signal probability (proportion of target to non-target stimuli and cuing) can be modified with AX-CPTs [43–45] that alter the classic CPT so subjects respond to a target (e.g., the letter “X”) only if it is preceded by a cue (e.g., the letter “A”). Along with allowing measurement of aspects of sustained attention and impulsivity [41], the AX-CPT differs from traditional CPTs for integrating features of working memory [46–48] and interference suppression [44]. In the context of SDT [3,4,49], AX-CPTs allow the quantification and distinction of sensitivity (d’) from response bias (β) with interference suppression [50]. The d’ measure provides a metric of separation between the signal and noise distributions, and quantifies the degree to which targets are successfully discriminated from non-targets given attention capacity [4]. The β variable measures response style, where low values indicate a willingness to accept many false alarms to keep hits high. This perspective on SDT has received widespread acceptance within cognitive neuroscience (e.g., [5,42,51,52]).

We used data from a set of healthy subjects, and assumed individual differences across the two tasks were stable to the degree that a definable mathematical structure was apparent between RPT and SDT variables. Such an assumption is warranted for RPT across temporal intervals of 1–2 weeks [11]. We first tested for mediation between variables (a statistical approach), and if none were found, then used an iterative modeling approach [53] to determine if any mathematical structure existed like a manifold, function, or boundary envelope. We hypothesized that a relationship would be observed between β, which describes the threshold one sets (i.e., a form of bias; [54,55]) between signal and noise distributions to determine the proportion of hits versus false alarms, and the {K, H} variables. We further hypothesized that no relationship would be observed between d’ from SDT, and {K, H}, given d’ relates to assessment of the proximity of noise and signal distributions [54].

## Methods

### Subjects

All subjects were recruited by advertisements from the New England region. Subject recruitment stopped after a set temporal window for recruitment, where target recruitment for healthy controls sought 50–100 subjects. This resulted in 77 subjects meeting criteria to be considered a healthy control in the Northwestern University and MGH Phenotype Genotype Project in Addiction and Mood Disorder (PGP; http://pgp.mgh.harvard.edu), who were not family members of a participant with cocaine dependence and polysubstance abuse, or a participant with major depressive disorder. Of these 77 subjects, 47 subjects were determined to have complete behavior, fMRI, and structural data for two reward/aversion tasks and a CPT task. In the analysis presented in the current manuscript, the data from the beauty keypress task and the CPT task are reported. Please see *Data Analysis* section for information about observation inclusion. For these subjects, the mean age was 31.0 years (SD 10.4), mean educational history was 15.6 ± 2.6 years, and subjects were 24/47 (51.1%) female, and 40/47 (85.1%) right-handed, with the following race identification: 36/47 European-American, 1/37 Native American, 6/47 African-American, and 4/47 Asian. All subjects underwent a clinical interview with a physician (psychiatrist) that included the Structured Clinical Interview for Diagnosis – Axis I (SCID-I/P; [56]). Race was determined by individual self-identification on a standardized form [57], and handedness via the Edinburgh Handedness Inventory [58]. Eligible subjects were age 18–55, without any current or lifetime DSM-IV Axis I disorder or major medical illness known to influence brain structure or function, including neurologic disease, or HIV and Hepatitis C as determined by assay. Medical illness was assessed via physician-led review of systems and physical exam. Female subjects were studied during their mid-follicular phase based upon self-reported menstrual history, with confirmation at the time of testing based on hormonal testing with a urine assay. All subjects were studied at normal or corrected normal vision.

### Ethics Statement

This study was approved by the Institutional Review Board (IRB) of Massachusetts General Hospital (i.e., Partners Human Research Committee, Partners Healthcare). All subjects signed written informed consent prior to participation, following procedures approved by the IRB. All experiments were conducted in accordance with the principles of the Declaration of Helsinki.

### Experimental Paradigms

#### a. Beauty Stimulus Set & Keypress Task

A scheduleless keypress task was used to determine each subject’s relative preference toward the ensemble of faces [11]. The ensemble of faces included the following experimental conditions: beautiful (models) and average (non-models) faces of both genders (i.e., beautiful female [BF], average female [AF], beautiful male [BM], and average male [AM]; [28]). The keypress procedure was implemented with MatLab software. This task captured the reward valuation attributed to each observed face, and quantified positive and negative preferences involving (i) decision-making regarding the valence of behavior, and (ii) judgments that determine the magnitude of approach and avoidance [11,28]. The objective was to determine how much effort each subject was willing to trade for viewing each facial expression compared to a default viewing time. Subjects were told that they would be exposed to a series of pictures that would change every eight seconds (the default valuation of 6 seconds + 2 second decision block; Figure 1a) if they pressed no keys. As published previously [11,28–35], each experimental stimulus would initially be presented for 0.2s, and be replaced by a fixation point for 1.8s, until the face came back at 2s and they could alter viewing time via keypressing or not. Keypressing to decrease or increase viewing time had a symmetric resistive function characterizing how more keypresses would result in ever smaller decrements or increases in viewing time. If they wanted a picture to disappear faster, they could alternate pressing one set of keys (#3 and #4 on the button box), whereas if they wanted a picture to stay longer on the screen, they could alternate pressing another set of keys (#1 and #2 on the button box). Subjects had a choice to do nothing (default condition), increase viewing time, decrease viewing time, or a combination of the two responses (Figure 1a). A “slider” was displayed to the left of each picture to indicate total viewing time. Subjects were informed that the task would last approximately 20 minutes, and that this length was independent of their behavior, as was their overall payment. The dependent measure of interest was the amount of work, in number of keypresses, which subjects traded for face viewtime. Keypress results could also be combined as total viewtime relative to the default baseline.

#### b. Attention Task

Subjects performed a continuous performance task (CPT; [42]) using visual rather than auditory stimuli [44]. In this task, subjects were required to respond to an “A” (target) following a “Q” (cue) after three intervening letters. This task added interference and divided attention load by, respectively, including false cues and/or false targets, and by intermingling a subset of QxyzA sequences within each other (e.g., QxQyAzA). Each letter was presented for 200ms and followed by a fixation point for 800ms. The task was administered as three “blocks” in the task, with each block lasting 60 seconds. Subjects responded to targets with a button press and did not respond to non-targets. Following a target detection framework [3–5], hits, missies, correct rejections, and false alarms were assessed and used in statistical analyses.

### Data Analysis

#### a. Computation of Operative Variables and Quality Assurance

##### Beauty Keypress Task

###### Descriptive Statistical Measures

Descriptive statistics were used to summarize keypress responses. We computed the mean intensity of K+ and K-location estimates (mean of the positive and negative responses respectively), and the Shannon entropy (uncertainty of making a choice; [27]) for the positive and negative keypress responses following procedures detailed elsewhere [10,11]. Please see Table 1 for summary statistics of K+, K-, H+ and H-. Observations with a mean keypress of 0 were excluded from the analysis since the logarithmic transformation of 0 does not exist. There were 7 observations with mean K-value of 0 (from 4 subjects) and 50 observations with a mean K+ value of 0 (from 26 subjects). Therefore, models with H+ as the dependent variable used 138 observations for the analysis, and models with H- as the dependent variable used 181 observations for the analysis. No exclusion of observations led to an exclusion of a participant.

**Table 1:** Descriptive Statistics for *β*, K and H

| Variable | N | Mean | S.D. | Min. | Max. |
| --- | --- | --- | --- | --- | --- |
| $\beta$ | 188 | 2.63 | 1.279 | 0.75 | 5.47 |
| $d'$ | 188 | 2.49 | 0.668 | 0.79 | 3.65 |
| K+ | 188 | 12.36 | 18.679 | 0.00 | 86.95 |
| K- | 188 | 10.94 | 7.957 | 0.00 | 29.30 |
| H+ | 188 | 1.72 | 1.626 | 0.00 | 4.24 |
| H- | 188 | 3.16 | 1.360 | 0.00 | 4.32 |

Legend: N (47 subjects x 4 observations per subject) is the total number of observations and S.D. is the standard deviation. The minimum and maximum values provide information on the observed range for the variable in the data.

###### Testing for Relative Preference Theory Structure

Location estimates (K+, K-) and Shannon entropy estimates (H+, H-) were evaluated to assess if graphical structure was observed in the form of power function with individual data that was consistent across subjects, and also observed as an envelop for group data with the same functional form as the individual data. These procedures are described elsewhere [10,11], and shown for one subject from the cohort and for group data from this cohort in Figure 1c.

###### Assessing recurrent relationships between location/dispersion measures

Power law scaling was assessed between variables by mathematical evaluation of a power law function fit to the observed graphical structure. If there was a recurrent structure confirmed across experimental conditions, then the similarities in graphical structure were assessed between (i) group data, and (ii) individual graphs. The graphical similarity was evaluated for each individual to determine if there was a similar mathematical form (albeit with different parameter fits) to the envelope from group data.

##### Attention Task

Signal detection estimates of β and d’ were computed after first computing the signal detection parameters for hits and false alarms [4,5]. Hits were computed as the number of correct responses for identifying targets to the total number of (true) targets. False alarms were calculated as the number of responses for non-targets to the total number of false targets and false cues. With these hit and false alarm rates, we then computed β (beta) and d’ (dprime) following standard SDT metrics (see Table 1) with the following Matlab code:

~~~
<MatLab code for computing d-prime and beta>
function [dprime, beta] = sdt(hit, fa)
z_hit = norminv(hit, 0, 1);
z_fa = norminv(fa, 0, 1);
dprime = z_hit - z_fa;
beta = normpdf(z_hit, 0, 1)/normpdf(z_fa, 0, 1);
~~~

#### b. Mediation Analysis

Kim and colleagues [10,11] found a significant relationship between K (independent variable) and H (dependent variable). To understand the role of β, we first examined whether H plays a mediating role between K and β, and found no significant relationship between H+ or H- and β. We then examined whether K plays a mediating role between β and H. Here, we found that while β has an insignificant effect on K+, its influence on K- is significant. Since the coefficients of β and K+/K- are both significant in a model that examines their impact on H+/H- (see Table 2), it seems that K partially mediates the relationship between β and H. We, therefore, decided to focus on a model that includes both K and β as explanatory variables and evaluate how they influence H.

**Table 2.** Estimated Coefficients for *β* and K from Different Functional Forms

| | | Logarithmic:<br>$H = a + b \cdot \log \beta + c \cdot \log K$ | Power Law:<br>$H = a * (\beta)^b * (K)^c$ | Power Law Additive:<br>$H = a + \beta^b + K^c$ |
| --- | --- | --- | --- | --- |
| **H+** | **Parameter** | **Estimate** | **Estimate** | **Estimate** |
| Intercept | a | 0.840**<br>[.52, 1.15] | 1.063**<br>[.86, 1.26] | -1.117**<br>[-1.41, -.82] |
| $\beta$ | b | 0.321**<br>[.04, .60] | 0.092*<br>[-.01, .20] | 0.239**<br>[.04, .43] |
| K+ | c | 0.667**<br>[.58, .75] | 0.319**<br>[.27, .37] | 0.349**<br>[.32, .37] |
| *RMSE* |  | 0.868 | 0.867 | 0.868 |
| *R* |  | 0.808 | 0.809 | 0.808 |
| **H-** | **Parameter** | **Estimate** | **Estimate** | **Estimate** |
| Intercept | a | 1.226**<br>[1.03, 1.43] | 1.522**<br>[1.37, 1.68] | -0.542**<br>[-.76, -.33] |
| $\beta$ | b | 0.196**<br>[.05, .34] | 0.044*<br>[.00, .09] | 0.141**<br>[0.01, 0.27] |
| K- | c | 0.951**<br>[.89, 1.01] | 0.337**<br>[.30, .37] | 0.443**<br>[.42, .46] |
| *RMSE* |  | 0.511 | 0.566 | 0.593 |
| *R* |  | 0.911 | 0.890 | 0.878 |

#### c. Assessment of Interactions between RPT and SDT Variables

We assessed the graphical interaction of {K, H, β} through iterative modeling [53], as done on prior occasions to determine if any mathematical structure existed like a manifold, function, or boundary envelope [10,11]. Given observation of two-simplex manifolds for the positive and negative components of the KH value function (Figure 2a-c), we checked for model stability, using random number generation to produce initial parameter estimates, and found no change in the estimated parameters. The fitting for these manifolds indicated H was a function of K and β in (a) a logarithmic relationship [H = log a + b·log β + c·log K; e.g., Figure 2a-c], (b) a multiplicative power law formulation [H = a·β
^b^·K
^c^], or (c) an additive power law formulation [H = a + β
^b^ + K
^c^; Table 2]. Fits for these three models were tabulated in Table 3, and computed in the following manner.

**Table 3.** Model Fits for H+ and H-Using Different Functional Forms

| Dependent Variable | Functional Form | RMSE | R |
| --- | --- | --- | --- |
| H + | Logarithmic | 0.8682 | 0.8079 |
| H - | Logarithmic | 0.5107 | 0.9112 |
| H + | Power multiplicative | 0.8667 | 0.8086 |
| H - | Power multiplicative | 0.5656 | 0.8899 |
| H + | Power additive | 0.8675 | 0.8082 |
| H - | Power additive | 0.5930 | 0.8782 |

Legend: Root Mean Square Error (RMSE) and R are both measures of model fit. R is the Coefficient of Determination and is the square root of the ratio of Regression Sum of Squares (SSR) to Total Sum of Squares (SST).

**Figure 2:**
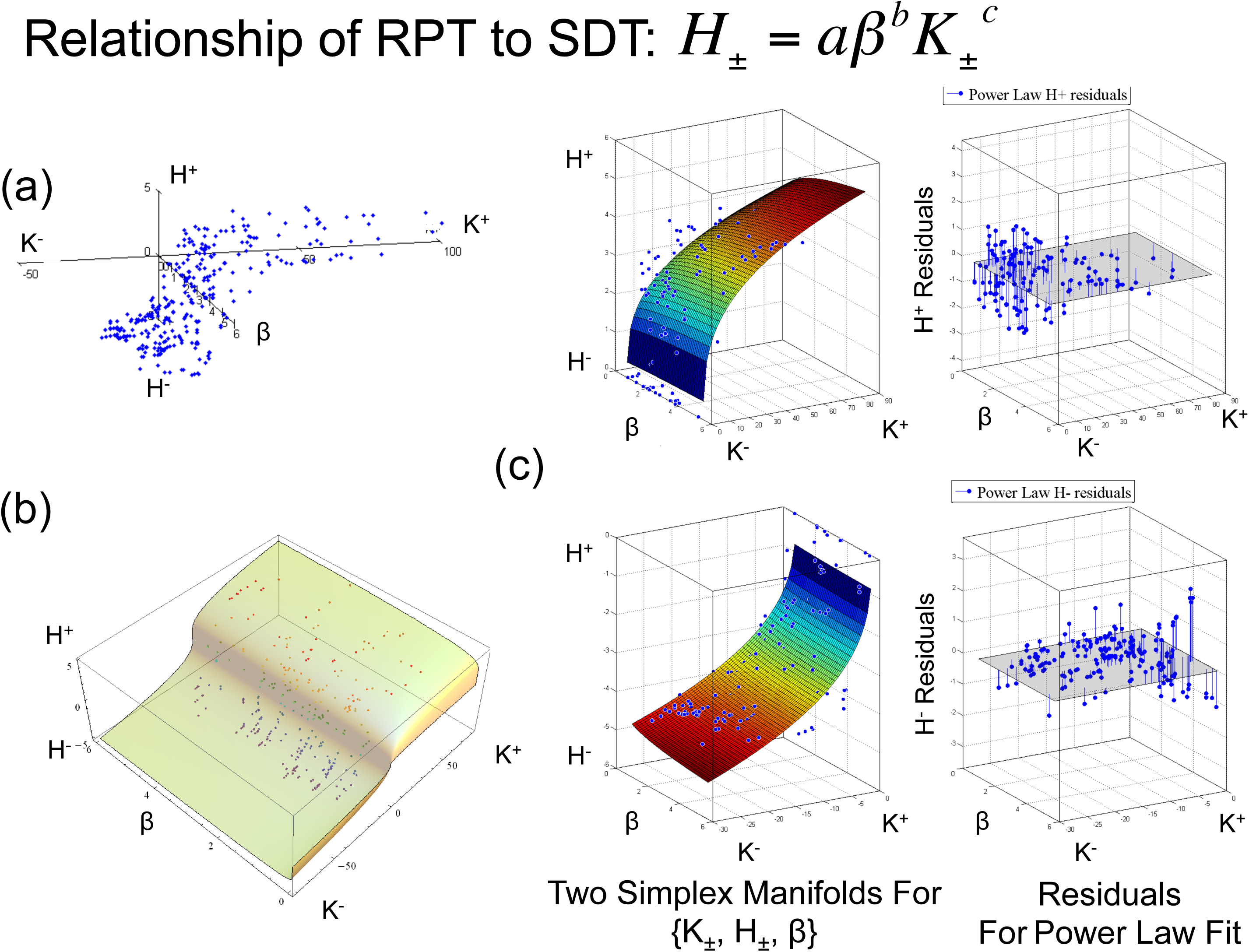
Iterative Modeling. (a) When K, H (in Figure 1b) include β as a z-axis, one observes a two-simplex manifold. (b) Fitting in Mathematica shows curvature along the β axis for approach and avoidance. (c) The twisting of the approach and avoidance manifolds is more obvious with colorized graphs in Matlab showing residuals of these graphs for approach (above right) and avoidance (below right). The approach manifold resembles a Cobb-Douglas graph.

To assess the strength of the fits for these three formulations of H α {K, β}, we computed Root Mean Square Error (RMSE) for each formulation to allow a comparison between fits. RMSE was computed as follows: 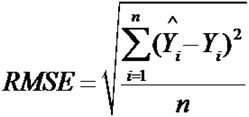, where 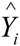is the predicted value of the dependent variable, *Y_i_* is the observed value of the dependent variable for observation *i* of *n*. We also computed R, where R is the Coefficient of Determination and is the square root of the ratio of Regression Sum of Squares (SSR) to Total Sum of Squares (SST). While SSR is the measure of the variation of the fitted regression values around the mean, SST is the measure of the variation of the observed values around the mean. Therefore, 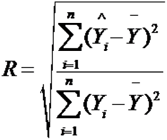, where 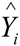is the predicted value of the dependent variable, *Y_i_* is the observed value of the dependent variable for observation *i* of *n* and, 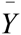is the average observed value of the dependent variables for all *n* observations.

In parallel with these assessments of functional fits between variables, we used the results of the power law estimation to get a better understanding of how β influences H+/H- and K influences β (Table 4). To partial out the effects of β on H+/H-, we computed the predicted values of H+/H- as follows:

**Table 4:** Assessment of Structure between RPT (Keypress) and SDT (Attention) Tasks

| Functional Form | $\beta = \alpha \cdot (H+)^{\gamma}$ | $\beta = \alpha \cdot (H-)^{\gamma}$ | $K+ = \alpha \cdot \beta^{\gamma}$ | $K- = \alpha \cdot \beta^{\gamma}$ |
| --- | --- | --- | --- | --- |
| $R$ | 0.03 | 0.04 | 0.08 | 0.04 |
| $RMSE$ | 1.329 | 1.251 | 20.018 | 7.764 |

Legend: Root Mean Square Error (RMSE) and R are both measures of model fit. R is the Coefficient of Determination and is the square root of the ratio of Regression Sum of Squares (SSR) to Total Sum of Squares (SST).

1. Use the respective estimated coefficients for β and K
2. Use the intercept estimated from the approach model as a common intercept for both approach and avoidance i.e., a = 1.522
3. Keep K constant for both K+ and K- at mean K+=12.36
4. Finally, vary β in small steps over the range [0,6] (please note the range of β in the data is slightly smaller at [0.75, 5.47]) to compute H+ and H-

This allowed a plotting of the predicted values of H+ and H- (y-axis) vs. β (x-axis), and assessment of their (i) ratio and (ii) difference as proxy measures of loss aversion (see Figure 3a).

**Figure 3:**
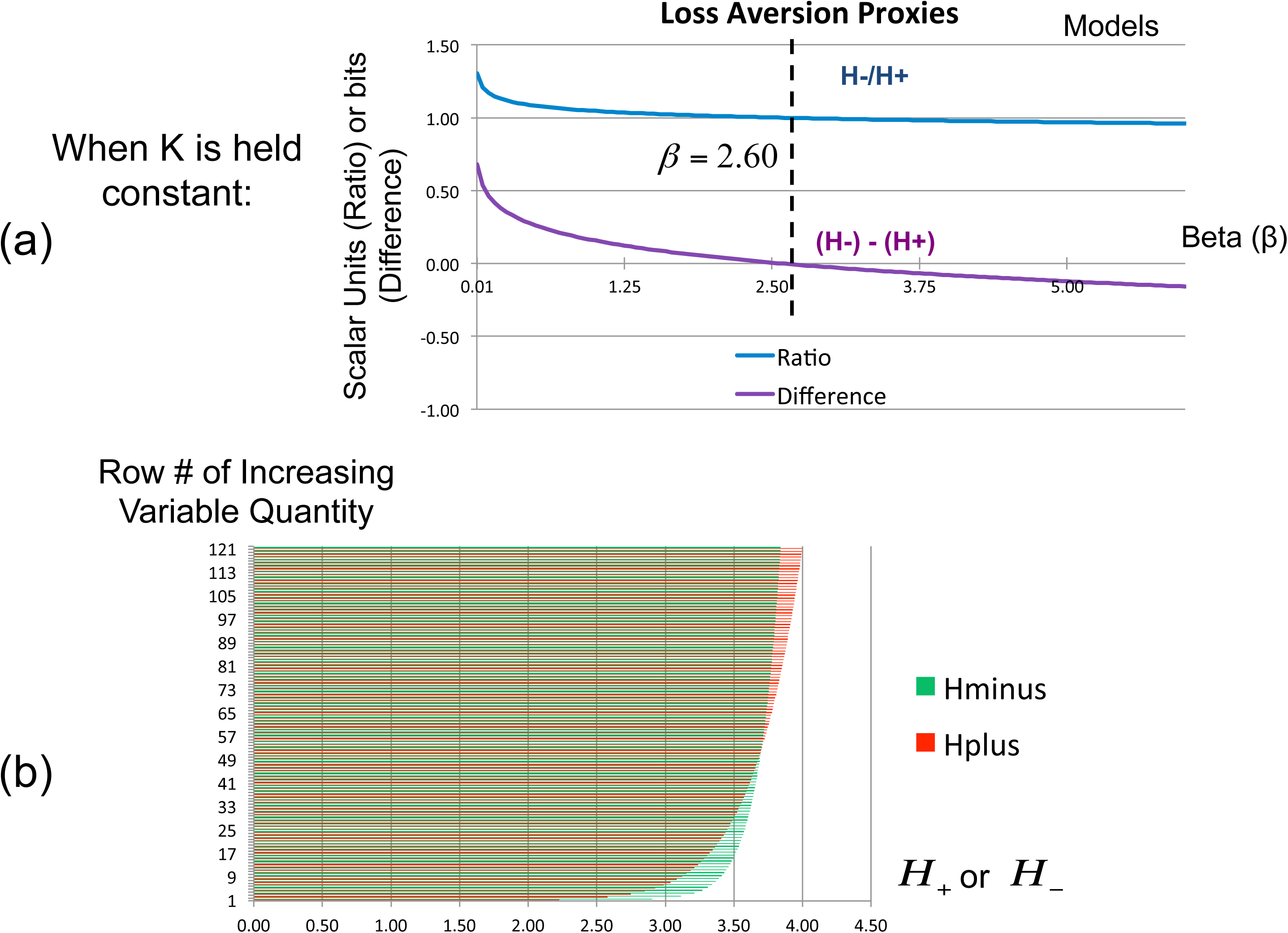
Evaluation of the Cobb-Douglas Function, and H± vs. β when K± is kept constant. (a) Using the ratio and difference of H- and H+ as proxies for loss aversion, graphed against β, we see both proxies cross mathematical inflection points (H-/H+ at 1, and (H-)-(H+) at 0) when β = 2.60. (b) When K is kept constant, and variables ordered by increasing quantity, there is a clear crossing of H+ and H- between line 49 and line 57.

To interpret this interaction, we further evaluated rank ordered data, whereby H+ and the absolute value of H- are ordered by iterative increases in β from values of 0.01 to 6.00, in increments of 0.05 (e.g., 0.01, 0.05, 0.10, 0.15…; see Figure 3b).

Robustness checks that included using the mean value of K-rather than that of K+, and using the intercept estimated from the avoidance model rather than the approach model did little to change the results.

To assess the potential quantitative interaction between RPT-based metrics of loss aversion and SDT-based measures of false alarms suggested in Figure 4a,b, we first computed loss aversion in the following way. In prospect theory, loss aversion has been defined as (i) the slope of the negative value/utility function (s-) compared to (ii) the slope of the positive value/utility function (s+), approximating the absolute value of s-/s+ (i.e., |s-/s+|; [9,59,60]). We applied a local definition of loss aversion [61–63] to the individual KH graphs (e.g., Figure 1c), wherein s- and s+ were computed by the integral of the curve-fit slope over the 10% of the curve closest to the inflection point or origin:

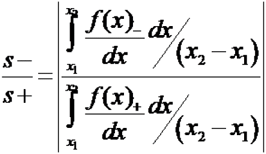

**Figure 4:**
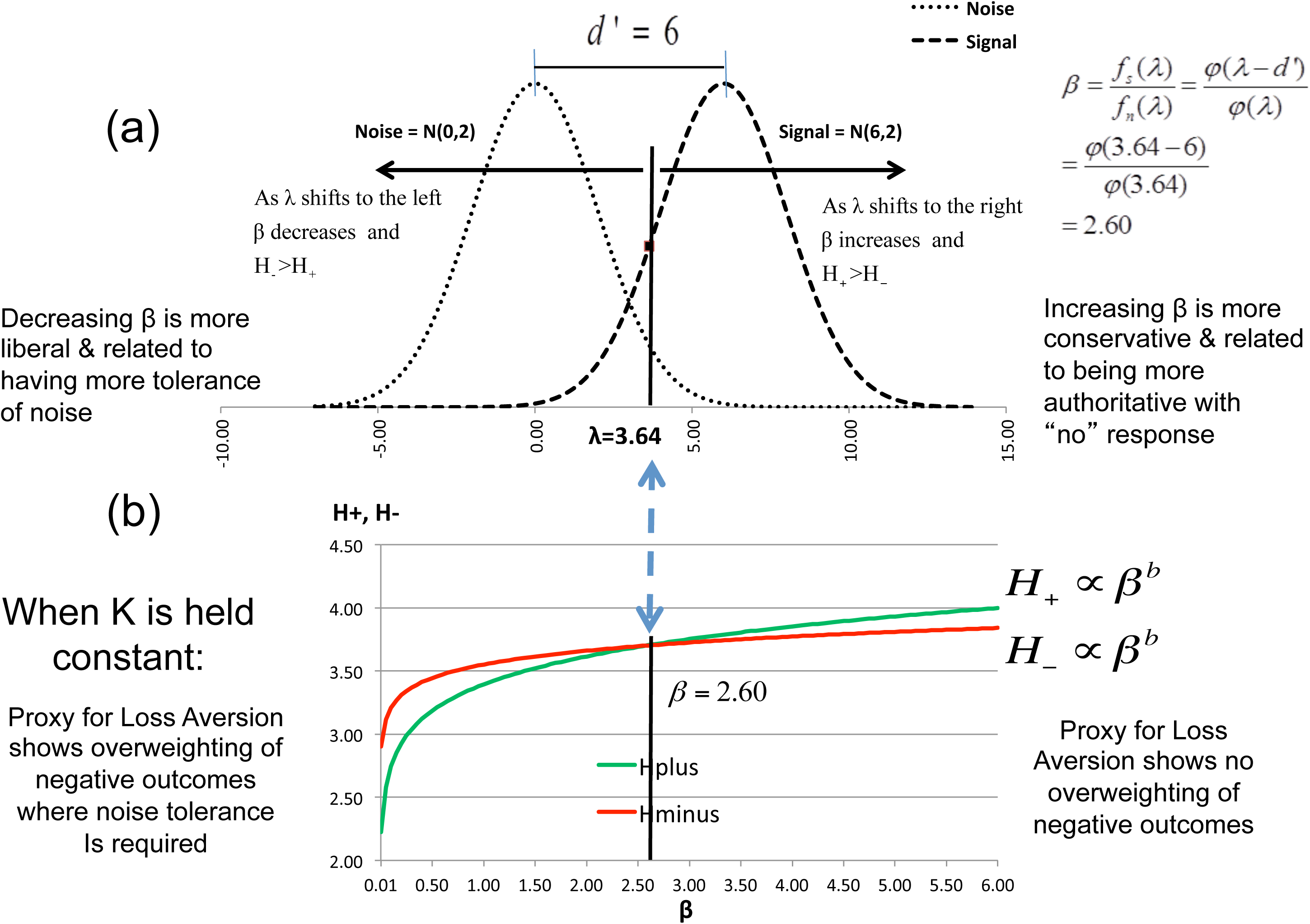
Interaction of {K, H, β}. (a) The signal detection schema and results are shown, where d’ = 6 and λ = 3.64, indicating β = 2.60 as the threshold where H+ vs. β and H- vs. β curves cross each other. (b) The crossing of curves at β = 2.60 shows how the loss aversion proxy is evident for β < 2.60, but absent (reflecting reward dependence) for β > 2.60.

We then used an absolute value of s-/s+ in a regression analysis with the SDT false alarm estimates for divided attention during the continuous performance task. Both linear regression and logarithmic regression (via log transformation of the x-axis) were performed (and reported) given potential skew in the loss aversion measures.

We used the threshold value of β observed in Figure 4a,b to determine the criterion parameter λ. Specifically, we used the illustration in Figure 4a,b to estimate the threshold criterion λ for an event where the distribution of noise is N(0,2) and that of the signal is N(6,2). Since the noise and signal have equal variances, we can calculate the parameter *d’* as the distance between the means of the two distributions = 6–0 = 6. *β* is calculated as 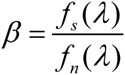, where *f_s_(λ)* and *f_s_(λ)* are the probability mass functions for signal and noise respectively. Hence, *β* is the ratio of the heights of the two distributions at the criterion, and using a standard normal table we can see that β = 2.60 when λ = 3.64

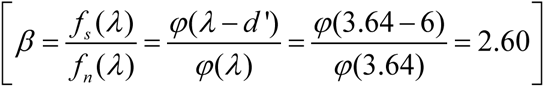

## Results

K, H, β, d’ data were compiled from 47 healthy subjects (Table 1).

### Assessment for Mediation

Given a known power law relationship between K and H [10,11], we first assessed if there was mediation between any three variables, and found none explaining more than 10% of the variance.

### Iterative Modeling {K, H, β}

*Graphical structure:* We assessed the graphical interaction of {K, H, β} through iterative modeling [53], and found two-simplex manifolds for the positive and negative components of the value function (e.g., Figure 2a-c), across multiple formulations with significant parameter fits (Table 2). To check for model stability, we used a range of initial parameter estimates and found no change in the results.

*Functional Form of Observed Structure*: We examined different functional forms and consistently found that K and β as explanatory variables explained 65–83% of the variance in H. Three functional forms were observed:

a. a logarithmic relationship [H = log a + b·log β + c·log K; Figure 2a-c],
b. a multiplicative power law formulation [H = a·β ^b^·K ^c^],
c. an additive power law formulation [H = a + β ^b^ + K ^c^].

Between the three functional forms, the multiplicative power law formulation [e.g., generalized as H_±_ = d ± a·(β ± f)
^b^-(K_±_ ± g)
^c^, where fitting parameters d, f, g = Ø with the current data] produced the lowest rank root mean square errors (RMSE) across both approach and avoidance fits (summarized in Table 3). This function is commonly referred to in economics as the Cobb-Douglas production function [64]. When fitting of the three formulations for {K, H, β} was tried with the constraint that the functions pass through the origin, all RMSE measures worsened, which contrasts with prospect theory [8,9] whereby all utility functions intersect the origin. We also verified that β was not significantly influenced by H+ or H- and confirmed the same (Table 4).

### Iterative Modeling {K, H, d’}

To complete the necessary negative control for specificity of the relationship found with {K, H, β}, we assessed the variables {K, H, d’} and observed no graphical structure between these variables.

### Exploratory Analyses of Findings

*Relationship of K-, K+, and β:* Other than a significant intercept, use of a power law estimation did not find β significantly influencing K+ or K- (Table 4).

*Relationship of H-, H+, and β:* When one partials out the effects of β on H-/H+, one observes that while the values of H- are higher than H+ for small values of β, H+ values increase at a faster rate than H- as β increases. Given β does not significantly affect K±, one can look at the ratio H-/H+ and the difference (H-) - (H+) as proxies for the measure of loss aversion, which traditionally is described as a ratio of slopes [8,9] although the difference model has been used with neuroimaging [65]. As a function of β, the ratio H- /H+ crosses the dimensionless value of 1 at a β of 2.6; similarly, the difference (H-) - (H+) crosses the value of 0 bits at the same β value (see Figure 3a).

To confirm the relationship between H+ and H-, we evaluated rank-ordered data (Figure 3b). When K is kept constant, and H-/H+ ordered by rank, there is a clear crossing of H- and H+ between lines 49 and 57 (Figure 3b).

These results are schematized in Figure 4a,b; namely, when one holds K+/K-constant, and evaluates the relation of β and H±, one observes that for β under a value of 2.6, H-is greater than H+, consistent with the concept of “loss aversion” in prospect theory [8,9]. In contrast, above β = 2.6, H+ is greater than H-which suggests greater sensitivity to reward seeking.

The relationship between graphs in Figures 4a,b suggests a positive correlation may exist between metrics of loss aversion from RPT and false alarms from SDT. We assessed this for a positive relationship and found that r(1,30) = 0.305, β < 0.048 as a linear regression, and r(1,30) = 0.353, β < 0.026 with log transformation of the x-axis. These results are consistent with the interactions shown in Figures 3a,b and 4a,b.

*Relationship of exponents for H = a·β ^b^·K ^c^:* For our data, b + c = 0.411 for approach, and b + c = 0.381 for avoidance. In Cobb-Douglas resource matching terms, this means that individuals show decreasing returns to scale or “inelasticity” in the form of reduced relative increases of H± despite substantial increases in (K±, β).

## Discussion

The relationship H α {K, β} means that equations modeling SDT [β = e
^(d’xC)^] [4,5] and RPT [H = a(K ± d)
^c^ ± f] [10,11] can form a function space in mathematical terms (Figure 2a-c). Two general implications arise from this finding. One implication relates to the concept of “loss aversion”, initially developed from prospect theory [8,9], but also observed with RPT [66]. Namely, loss aversion in the intrinsic motivation framework of RPT appears to be dependent on how individuals set the threshold for β in SDT. The more β decreases, consistent with increasing tolerance of noise in the form of more false alarms, the more one observes a relative prioritization of H-relative to H+, which in the context of holding K± constant, means the slope of the avoidance curve (i.e., K-H-) is steeper than the slope of the approach curve (i.e., K+H+). These data argue that “loss aversion” may be a property of decision-making when individuals must be more tolerant of lower signal-to-noise, and indeed a statistically significant positive interaction is observed between loss aversion estimates and false alarms. When there is high signal-to-noise, H+ is prioritized relative to H-, and individuals are mainly responsive to gains. One observes the exponent b for β in equation H = a·β
^b^·K
^c^ shows a proportion of 2:1 for approach:avoidance, which is not the case for the exponent c (Table 2), indicating the crossing of curves in Figures 3b and 4b are due to attention-related effects. This is intriguing in the context that β in SDT is a likelihood measure [3,4], and hence relates to uncertainty, further supporting its relevance to the relationship between approach and avoidance. Altogether, signal detection appears to be important for the emergence of loss aversion.

A second implication arises when one considers that the multiplicative power law observed with our data (i.e., H = a·β
^b^·K
^c^) has the same format as the Cobb-Douglas production function observed in economics [64]. In economics, the Cobb-Douglas function has been broadly used, in applications ranging from matching theory for describing mutually beneficial relationships [67], to the integration of real money balances into the production function to model the private domestic sector of the US economy from 1929–1967 [68]. An important feature of Cobb-Douglas is that the power law exponents together determine the relationship between the input variables (i.e., K±, β) and the output variable (H±) as a type of resource matching operation [64]. For instance, the output variable doubles when the inputs double and the exponents b + c = 1. When b + c > 1, the output variable (H) will show increasing returns to scale or higher “elasticity” in terms of small changes in the independent variables (K±, β), leading to larger changes in the dependent variable (H±). In contrast, if b + c < 1, the output variable will show decreasing returns to scale or “inelasticity” as reduced relative increases of H± despite substantial increases in (K±, β). For our data, b + c << 1 for both approach and avoidance, consistent with a control function defining capacity constraints [67] to mental processing for attention and reward/aversion.

Our data shows that a relationship exists between quantitative formulations of reward/aversion and attention. The relationship H α {K, β} can be expressed as a multiplicative power law (i.e., Cobb-Douglas function) that in economics reflects resource matching operations of potential relevance for constraining the relationship between attention and reward/aversion in behavior. This relationship underscores why concerns have arisen about the potential interaction of attention with reward/aversion in psychology and neuroscience [19], and raises at least two issues. First, when reward/aversion and attention tasks are performed in isolation, the results may need to carry a caveat about the other function not being controlled in the experiment. Second, the relationship observed between SDT and RPT in Figure 2c allows iterative modeling of the interaction between these variables as shown in Figures 3a,b and 4a,b. This interaction suggests that an engineering-based approach to behavioral science may be possible that allows variables across many behavioral domains to be connected and schematized as currently done for biochemical pathways. Such an interaction between variables resembles what is observed for the gas laws in thermodynamics, wherein one variable can be held constant, allowing the interaction of the other two to be explicitly modeled. This type of interaction is mechanistic, and explicitly allows for inference. If confirmed across other studies, such findings would imply that behavior modeling could be mechanistic, potentially to a similar degree as in chemistry. For this to occur, more work is needed given that domains such as attention encompass a broad field (e.g., [69]), and we have only shown how endogenous (vs. exogenous), overt (vs. covert) attention processes involving cognitive load (vs. perceptual load) is related with an RPT model of reward/aversion.

## Acknowledgments

Support for this work was provided by grants to Dr. Breiter (#14118, 026002, 026104, 027804) from NIDA, and grants (DABK39–03–0098 & DABK39–03-C-0098; The Phenotype Genotype Project in Addiction and Mood Disorder) from the ONDCP - CTAC, Washington, D.C. Further support was provided by a grant to A.J.B. (#052368) from NINDS. Additional support for this work was provided by the Warren Wright Adolescent Center at Northwestern Memorial Hospital. We thank Daniel Stern for his research around Cobb-Douglas functions, and James Reilly for his suggestion to check the correlation of loss aversion with false alarms. Dr. Mortensen had a fundamental impact on this work (being a dominant force interpreting results), and the authors accordingly dedicate this manuscript to his memory.

